# The Zika Virus Persistently Infects Retinal Pigment Epithelial Cells, Eliciting Limited Cytopathic Effects and Loss of Epithelial Characteristics Consistent with Epithelial-Mesenchymal Transition

**DOI:** 10.1101/2025.07.19.665681

**Authors:** Daed EL Safadi, Alexandre Mokhtari, Grégorie Lebeau, Wildriss Viranaicken, Pascale Krejbich-Trotot

**Author notes:** Correspondence (P.K-T.).

## Abstract

The Zika virus (ZIKV), a pathogenic member of the orthoflavivirus family, is raising serious health concerns worldwide. Like Dengue (DENV) and Chikungunya (CHIKV) viruses, it is one of the arboviruses that poses an emerging threat to areas where its main vectors, *Aedes* mosquitoes, proliferate, well beyond tropical and subtropical regions. Although often asymptomatic or mild, ZIKV infection has been responsible for a worrying increase in serious congenital syndromes, including microcephaly. The ability of ZIKV to be transmitted sexually and its long persistence in body fluids suggests its incomplete clearance in particular tissues, linked to recurrent infection. Among its clinical presentations, ZIKV infection has been associated with ocular complications, including maculopathy, retinopathy, uveitis, and optic neuropathy, which can lead to lasting visual impairment. The blood-retinal barrier (BRB), primarily composed of retinal pigment epithelium (RPE) and endothelial cells, plays a crucial role in shielding the retina from pathogens. Its disruption has been linked to viral retinal infections. In vitro monitoring of infection on hTERT RPE-1 cells revealed an ability of ZIKV to persist for up to 30 days in nearly 10% of the cells. This prolonged infection was marked by low cytopathic effects but notable morphological changes throughout the cell layer, suggestive of an epithelial-mesenchymal transition (EMT). Long-lasting viral replication and production was associated with reduced expression of epithelial genes and increased expression of certain mesenchymal genes, suggesting that integrity of the RPE layer may be compromised. These results indicate that viral persistence and phenotypical transition observed in vitro in RPE cells could provide clues to understanding the late onset of ocular pathophysiological manifestations in Zika virus-related diseases.

## 1. Introduction

Zika virus (ZIKV) belongs to the *Orthoflavivirus* genus and represents a significant public health threat. This arbovirus is primarily transmitted by mosquitoes of the *Aedes* genus. In addition to mosquito-borne transmission, ZIKV can also spread through horizontal transmission via sexual contact [1], and vertical transmission from an infected mother to her fetus [2]. ZIKV infection is often asymptomatic, and when symptoms occur, they are generally mild and transient, including fever, headache, rash, nausea, myalgia, and arthralgia [3]. However, infection can lead to severe complications and potentially fatal conditions. In particular, the outbreaks of 2013 in French Polynesia and 2015 in Brazil revealed the possibility of nervous system damages in adults, with an increased incidence of Guillain-Barré syndrome [4,5]. They were also characterized by the onset of serious and alarming congenital anomalies in children whose mothers had been infected during pregnancy. The combination of clinical manifestations, including microcephaly, has led to the definition of a congenital Zika syndrome (CZS) [6].

It has also been reported that ZIKV can cause ocular lesions and ophthalmic complications [7,8]. Remarkably, up to 50% of infants with Zika-related microcephaly present ocular abnormalities [9]. While these complications are most pronounced in newborns, it has been documented that ZIKV infection can lead to visual impairment in adults. Among infected adults, conjunctivitis occurs in 10 to 15% of cases [10]. Surprisingly, some uveitis has been reported weeks after the initial infection with detection of viral RNA in the aqueous humor [11]. Maculopathy has been observed in a case report [12]. In certain cases, visual impairment may persist for several months [13]. In addition, experimental models have confirmed ZIKV’s tropism for the eye and its ability to induce long-lasting lesions and retinopathies in mice [14,15].

Ophthalmic damage due to viral infections indicates blood-retinal barrier (BRB) fragility and potentially disruption of the retinal epithelium and/or endothelium [16,17]. The BRB, which extends from the blood-brain barrier (BBB), separates the eye’s internal environment from the systemic circulation. It consists of two components: an inner barrier formed by retinal endothelial cells and an outer barrier composed of the retinal pigment epithelium (RPE) layer, Bruch’s membrane, and the choriocapillaris. The external BRB, supported by tight junctions between RPE cells, protects the neuroretina from blood-borne pathogens, including viruses, serving as the first line of defense against pathogens in the retina [18].This barrier also contributes to innate and adaptive immunity. Disruption of the RPE can provide an entry route for viruses that cross the fenestrated choroidal capillaries to infiltrate retinal tissue [19]. The integrity of the BRB is crucial for healthy vision, and its impairment can lead to various retinal diseases. Several studies have shown that the cell components of the BRB, such as retinal epithelial and endothelial cells, are permissive to ZIKV and contribute to the inflammatory response [20–22].

However, the exact mechanisms behind the development of ocular complications and the fragilization of the BRB following ZIKV infection remain unclear. In order to explore the mechanisms behind this pathophysiological process, we characterized the infection of hTERT RPE-1 cells by a clinical isolate of the Asian epidemic strain of ZIKV (PF-25013-18). Interestingly, we found that hTERT RPE-1 cells were permissive to ZIKV and that infection could be maintained in vitro for more than 30 days. Analysis of cell death markers over time indicated that infection elicited a moderate cytopathic effect that did not compromise cell culture continuity. However, microscopic observation of the persistently infected hTERT-RPE monolayer suggested a morphological change, with cells taking a more elongated, fibroblast-like shape. Further characterization of this viral-induced phenotype by gene expression analysis showed that persistent infection led to a decrease in the expression of some epithelial markers (E-cadherin, Claudin-1) followed by an increase in the expression of two mesenchymal markers (fibronectin, TGF-β). These findings suggest that a potential epithelial-mesenchymal transition (EMT) could occur in chronically infected RPE and constitute the early steps of pathological consequences of viral infection in the eye.

## 2. Materials and methods

### 2.1. Viruses, cells and reagent

ZIKV is a clinical isolate of the Asian lineage, PF-25013-18 (ZIKV-PF13), obtained during the French Polynesian outbreak and previously described [23].

Immortalized retinal pigment epithelial cells (hTERT RPE-1), obtained by overexpressing the catalytic subunit of telomerase, were purchased from ATCC (CRL-4000). The cells were cultured in DMEM (Dulbecco’s Modified Eagle Medium from Gibco/Invitrogen, Carlsbad, CA, USA) supplemented with 10% heat-inactivated Fetal Bovine Serum (FBS Invitrogen), 1mmol/L sodium pyruvate, 2mmol/L L-Glutamine, 0.1mg/mL streptomycin, 100 U/mL penicillin, 0.5µg/mL fungizone and 50 µg/ml hygromycin B (PAN-Biotech, Aidenbach, Germany), as specified by the supplier for the selection pressure. For the infection experiments, hTERT RPE-1 cells were gradually switched to 1% FBS medium in order to accentuate the properties of a confluent cell layer exhibiting epithelial characteristics, as recommended in the literature [24].

### 2.2. Lactate dehydrogenase assay (LDH)

The hTERT RPE-1 cells were plated in a 24-well plate with seeding density corresponding to 100,000 cells per well. 24 hours later, cells were infected with ZIKV at different multiplicities of infection (MOI 0.1,1, and 5). The infection was stopped at different time points and the culture supernatants were collected. DMEM–dimethyl sulfoxide (DMSO) 10% was used as a positive death inductor. Following viral deactivation under UVs, the LDH assay was performed using the CytoTox 96® Non-Radioactive Cytotoxicity Assay (Promega), following the manufacturer’s guidelines. Briefly, 50 µL of LDH was added to 50 µL of LDH substrate and incubated at 37°C for 20 min. The absorbance was then read at 490 nm using a FLUOstar® Omega (BMG LABTECH, Offenburg, Germany).

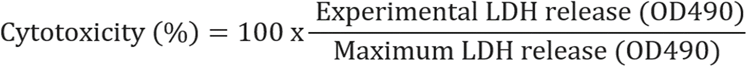

### 2.3. Flow cytometry and antibodies

Cells were harvested at different time points of infection using trypsin, fixed for 10 minutes using paraformaldehyde in PBS (PFA 3.7%), and permeabilized using Triton X-100 at 0.15% for 5 minutes. Immunostaining of ZIVE protein was done with the 4G2 antibody followed by Alexa Fluor 488 IgG secondary antibody. Fluorescent cells were quantified using a CytoFLEX flow cytometer (Beckman Coulter, Villepinte, France).

### 2.4. Immunofluorescence imaging of infection and cell death by apoptosis

hTERT RPE-1 cells were grown and infected on glass coverslips. The cells were then fixed with 3.7% formaldehyde in PBS at room temperature for 10 minutes. After fixing, the cells were permeabilized with 0.1% Triton X-100 in PBS for 5 minutes. The coverslips were incubated with primary antibodies : the mouse 4G2 (anti-panflavivirus E protein) from RD-Biotech (Besançon, France) was used to detect infected cells and the rabbit anti-BAX (#2772) from Cell Signalling Technology (Ozyme, Saint-Cyr-l’École, France) was used to detect apoptotic cells. Antibodies were diluted at a 1:1000 in 1× PBS with 1% BSA and added to coverslips for 1 hour, followed by incubation with Alexa Fluor 488 or 594 IgG secondary antibodies (1:1000, Invitrogen) for 1 hour. Nuclear morphology was visualized using DAPI staining. The coverslips were then mounted with VECTASHIELD® (Clinisciences, Nanterre, France). Fluorescence was observed using a Nikon Eclipse E2000-U microscope, and images were captured and processed with a Hamamatsu ORCA2 ER camera and NIS-Element AR imaging software (Nikon, Tokyo, Japan).

### 2.5. Measurement of virus production by plaque forming unit assay (PFU)

To quantify the release of infectious viral particles, plaque-forming unit (PFU) assays were performed. Vero cells were seeded the day before in 24-well plates at a density of 7×10^4 cells per well. The following day, cells were infected with 0.1 mL of ten-fold serial dilutions of the infected RPE cell supernatants. After a 2-hour incubation at 37°C, 0.2 mL of culture medium containing 5% fetal bovine serum (FBS) and 1% carboxymethylcellulose (Sigma-Aldrich, Saint-Quentin-Fallavier, France) was added, and Vero cells were incubated for an additional 4 days at 37°C. Cells were then fixed in 3.7% paraformaldehyde (PFA) and stained with 0.5% crystal violet solution in 20% ethanol. Plaques were counted and reported as plaque-forming units per milliliter (PFU/mL).

### 2.6. RNA extraction, primers and qRT-PCR

Total RNA was isolated from the RPE cell lysates using the RNeasy Plus Mini Kit (QIAGEN, cat. 74136, Hilden, Germany). Complementary DNA (cDNA) was generated via reverse transcription with random primers (Invitrogen, ref. 58875, Thermo Fisher, Waltham, MA, USA) and the M-MLV reverse transcriptase enzyme (Promega, ref. M1708, Madison, WI, USA) at 42°C for 60 minutes. The resulting cDNA was then analyzed by quantitative polymerase chain reaction (qPCR) using the CFX96 Connect™ Real-Time Detection System (Bio-Rad, Hercules, CA, USA). Amplification was performed with ABsolute™ 2X qPCR MasterMix (Thermo Fisher, ref. AB1163A, Waltham, MA, USA). For each sample, the threshold cycle (Ct) value was determined during the exponential amplification phase using Bio-Rad CFX Manager 3.1 software (Bio-Rad, Hercules, CA, USA). The qPCR results were analyzed using the ΔΔCt method (Livak and Schmittgen, 2001) and normalized to the expression of the housekeeping gene ribosomal protein lateral stalk subunit P0 (RPLP0). To test if EMT occurred during the timeline of infection, epithelial markers tested were human E-cadherin encoded by CDH1 gene and Claudin-1 encoded by CLDN1 gene and for mesenchymal markers, Fibronectin encoded by FN1 gene and TGF encoded by TGFB1 gene. Primers sequences were:

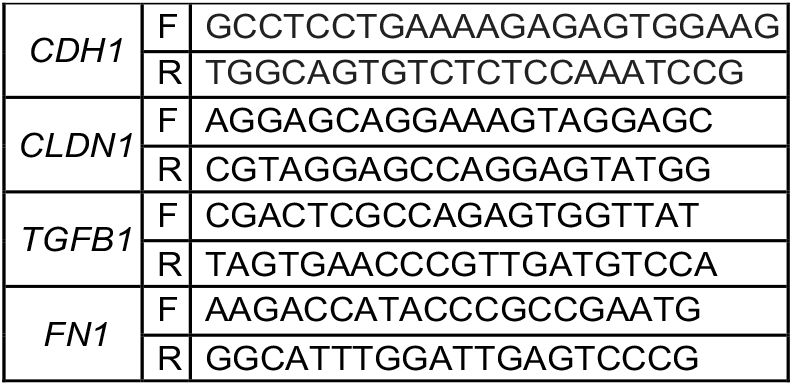

### 2.7. Statistical analysis

All values are expressed as mean ± SD of at least three independent experiments. Comparisons between infected and uninfected cells were analyzed using ordinary one-way ANOVA. Values of p < 0.05 were considered statistically significant. Significance levels are indicated in the figure legends as follows: ns = not significant, * p < 0.05; ** p < 0.001; *** p < 0.0002; **** p < 0.0001. All statistical tests were performed using GraphPad Prism version 9 software (GraphPad Software, San Diego, CA, USA, available online: http://www.graphpad.xn--com-9o0a).

## 3. Results

### 3.1. hTERT RPE-1 cells are permissive to ZIKV

RPE cells were infected with ZIKV-PF13 at MOIs of 0.1, 1, and 5 for 24, 48, or 72 hours. Infection rates were measured by flow cytometry and by microscopy imaging. Cytopathic effects were evaluated by measuring the catalytic activity of LDH released in the cell supernatant and by immunofluorescence imaging of the apoptotic marker Bax.

We confirmed that hTERT RPE-1 cells are susceptible to ZIKV **(Figure 1A)**, supporting previous studies on other RPE from different origins [22,25,26]. The cytopathic effects induced during the ZIKV infection time course were limited, with a slight increase over time reaching 25% of infected cells presenting BAX localized to mitochondria, 72 hours post infection (h.p.i) (**Figure 1B and 1C**). Interestingly, infection seems to peak at 48 h.p.i. At 72 h.p.i, regardless of the MOI, the percentage of infection was found around 15%. Based on these results, we set the MOI at 1 and decided to carry on the kinetics of infection over longer times.

**Figure 1:**
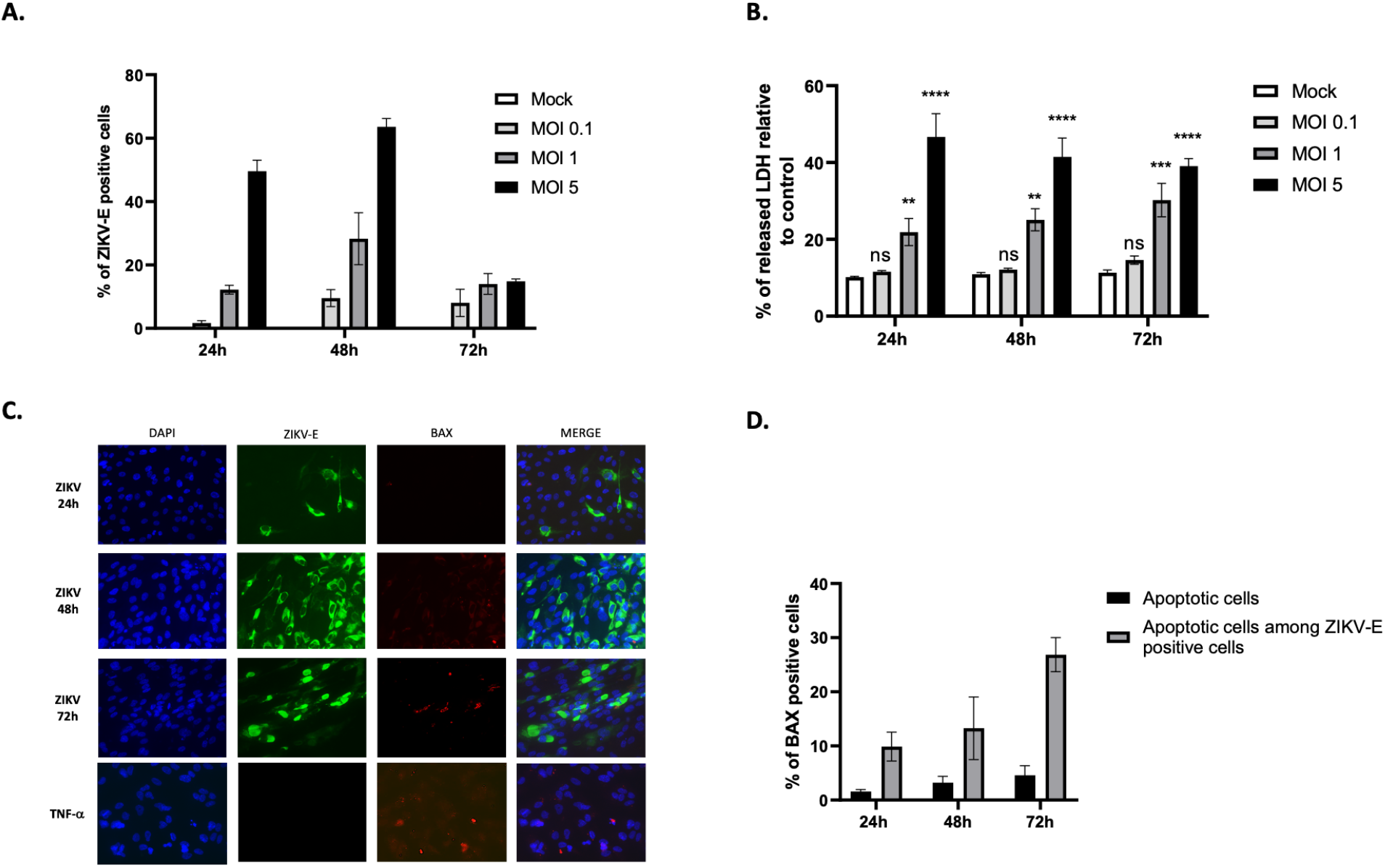
RPE cells are permissive to ZIKV with limited cytopathic effects. **(A)** RPE cells were infected with ZIKV at MOIs of 0.1, 1, and 5 for 24, 48, and 72 hours, and infection was assessed by flow cytometry. **(B)** Cytopathic effects related to cell necrosis were assessed at different time points and MOIs by measuring LDH release in cell culture supernatants. **(C)** Cells infected with ZIKV at MOI 1 or treated with TNF-a (apoptotic positive control) were immunostained for ZIKV E (green), BAX (red). Nuclei were stained with DAPI (blue). (**D**) The percentage of apoptotic cells, presenting mitochondrial BAX detection, in total and among ZIKVE-stained cells was estimated based on counting a minimum of 3 microscopic fields. Values represent the means and standard deviations of three independent experiments. \*\*\*\**p*<0.0001,\*\**p*<0.01

### 3.2. ZIKV persists in RPE cells for up to 30 days

Given the mild infection rates and limited cytopathic effects, we decided to follow the cell and virus fate by maintaining infected hTERT RPE-1 cells in culture for longer periods. RPE cells were cultivated in 1% FBS to maximize the epithelial characteristics of the confluent cell layer, and the medium was renewed every week. Confluent cells were infected with ZIKV-PF13 at MOI = 1. We monitored the infection at days 3, 7, 14, 21 and 30 using flow cytometry. We also regularly proceeded to microscopic imaging of the cell culture.

We observed that RPE cells infected with ZIKV and expressing the E protein were detectable for up to 30 days **(Figures 2A and 2C)**. Infection peaks seem to punctuate the kinetics, with a maximum of about 25% of cells immunolabeled for ZIKV E, observed at 48 h.p.i. and 7 days post infection (d.p.i.). The profile is interesting, alternating phases of decrease in the number of infected cells (to 2-3%) and phases of recovery up to 25%. Microscopic observation and quantifying the number of infected cells that exhibit mitochondrial BAX suggest an apoptosis rate of approximately 15–20% in ZIKV-E-positive cells. This value is constant over time, including 30 d.p.i (**Figure 2B**). RPE cells grown over time with a basal level of persistently infected cells by ZIKV, appear to maintain a compromise between cell death and cell growth, throughout culture.

**Figure 2:**
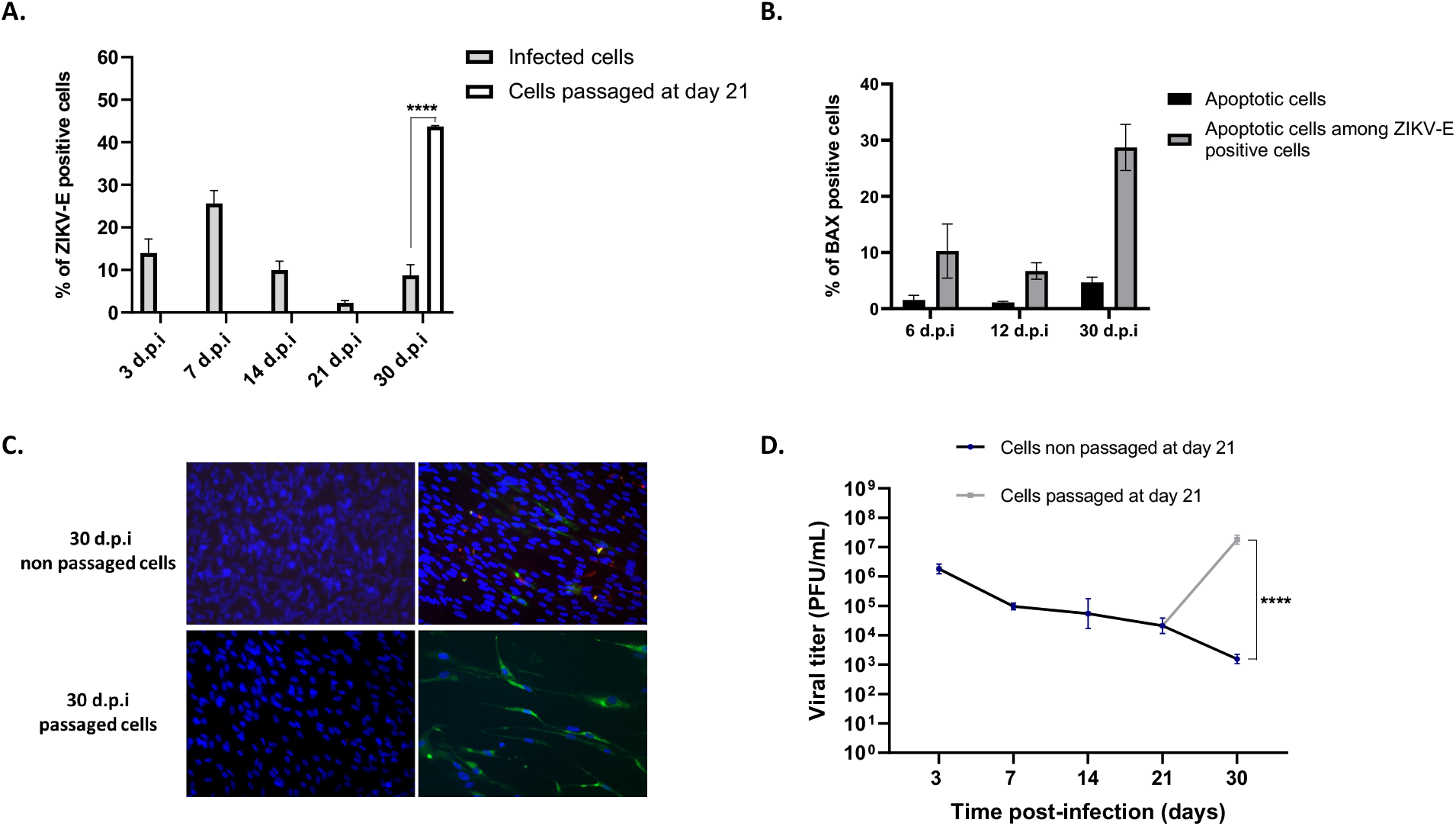
ZIKV persists in RPE cells. **(A)** Cells were infected at MOI 1 for 3, 7, 14, 21, and 30 days with or without passage at day 21. The percentage of ZIKV-E positive cells was measured by flow cytometry. **(B)** The percentage of apoptotic cells was estimated by immunodetection of BAX localization in mitochondria and based on counting at least 3 microscopic fields. **(C)** Immunofluorescence imaging of RPE infection at 30 d.p.i with ZIKV-E labeling (green), BAX labeling (red), and DAPI nuclei staining (blue). **(D)** Quantification by plaque forming unit assay of infectious ZIKV particles present in the cell culture supernatants during the 30-day kinetic study. Values represent the mean and standard deviations of three independent experiments. \*\*\*\**p*<0.0001

To ensure that ongoing infection was associated with the production of infectious particles, we performed PFU assays to determine the titer of the virus in cell culture supernatants, at different infection time points. We showed that cells continued to produce infectious viral particles up to 30 d.p.i, with a relatively stable titer of around 5.10^4^/10^5^ pfu.mL^-1^ estimated at 7, 14 and 21 d.p.i. (**Figure 2D**).

It is interesting to note that mock-infected cells were maintained in a confluent culture for 3 to 4 weeks without any problem, confirming the capabilities of RPE cells to form a confluent monolayer of quiescent cells [27]. We then wanted to compare the behavior of cells maintained in culture with or without passage reactivating their growth. At 21 d.p.i., the persistently infected cells were trypsinized and seeded in new plates, to the third of their initial cell density. This resulted in a significant increase in the percentage of ZIKV-infected cells 10 days later, at 30 d.p.i., rising from 10% to approximately 45% after cell passage (**Figure 2A and 2C**). Furthermore, the production of infectious viral particles by RPE cells was significantly higher at 30 d.p.i. following passage at day 21 compared to non-passaged cells, with an increase of approximately 4 logs, as shown in **Figure 2D**.

### 3.3. Cells persistently infected with ZIKV undergo epithelial-mesenchymal transition

EMT is a crucial physiological process for normal embryonic development and tissue regeneration, such as wound healing. However, under pathological conditions, EMT can become dysregulated, leading to tissue fibrosis and supporting cancer metastasis [28]. EMT correlates with a decrease in expression of E-cadherin, a protein that maintains adherens junctions between epithelial cells [29]. As cells transition to a mesenchymal state, they exhibit an increased expression of markers like N-cadherin and vimentin, along with extracellular matrix (ECM) proteins such as fibronectin and collagen [30]. Transforming growth factor-beta (TGFβ) is a major regulator and one of the most studied contributors to this process, increasing cell capacities to migrate and proliferate [31]. These factors involved in EMT are illustrated in **Figure 3A**. In some cases, EMT may be incomplete, referred to as partial EMT, where epithelial markers are not fully lost and mesenchymal markers are not fully gained. The reorganization of the cytoskeleton observed following EMT results with an elongated, spindle-shaped morphology, fibroblast-like, characteristic of mesenchymal cells [32].

**Figure 3:**
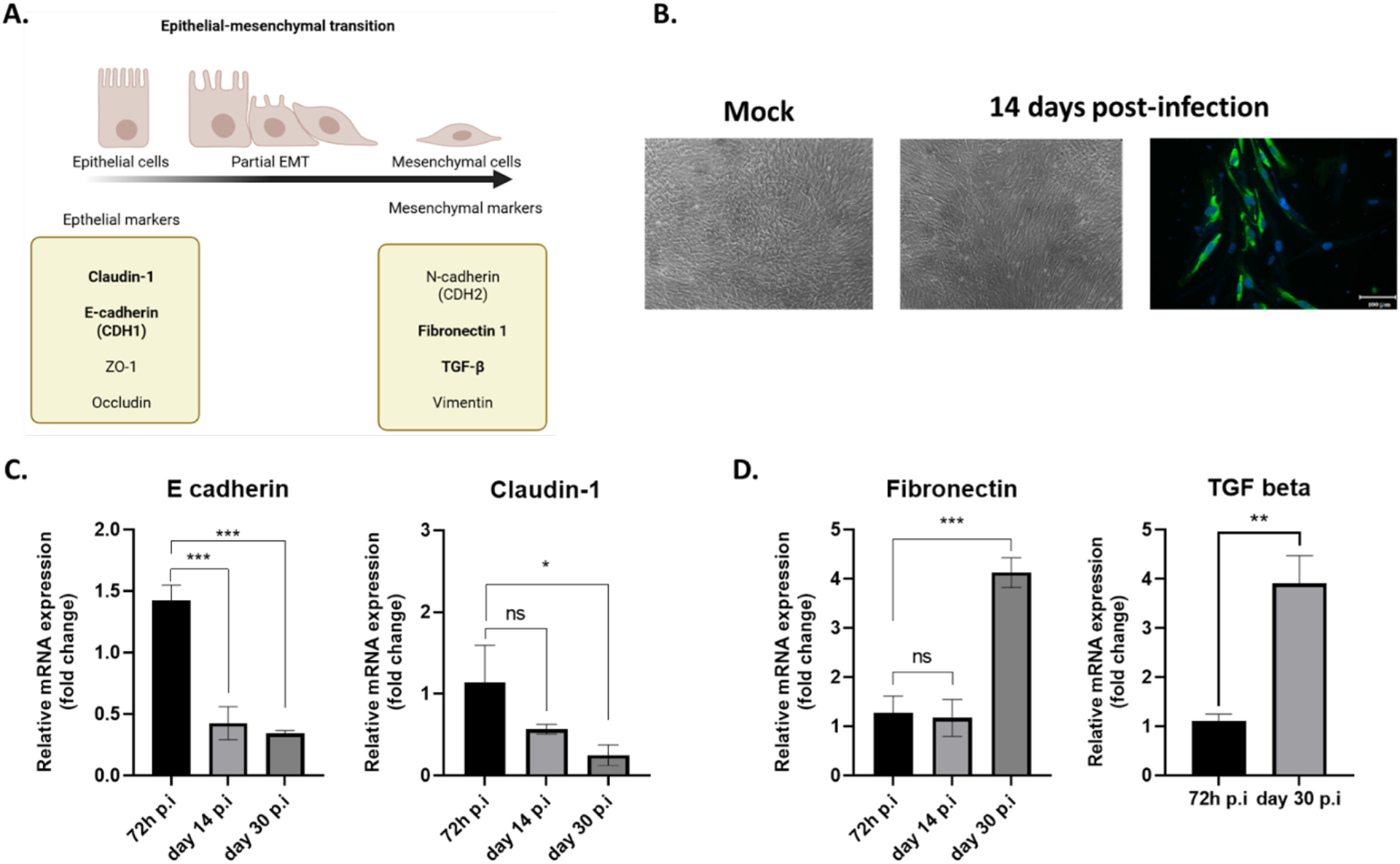
ZIKV persistently infected cells undergo partial EMT. **(A)** Diagram depicting the epithelial-mesenchymal transition, highlighting key markers involved. **(B)** Control cells and ZIKV-infected cells at 14 d.p.i were visualized by microscopy, showing that infected cells exhibit morphological changes, becoming elongated. The panel on the right shows infected cells immunostained for ZIKV E. Gene expression of markers involved in the EMT process was measured by qRT-PCR at 72 hours, 14, and 30 days post-infection. The expression of epithelial markers is shown in **(C)**, and mesenchymal markers in **(D)**. *** *p* < 0.001, ** *p* < 0.01

When following the long term cultures of RPE under the microscope, we observed that ZIKV-persistently infected cells changed their morphology to a fusiform shape, as revealed by comparisons of cell culture pictures (**Figure 3B**). This led us to hypothesize that cells undergo EMT. Cultivated RPEs have been shown to retain the reprogramming capacity to evolve along a continuum between polarized epithelial and mesenchymal cells [30]. We therefore explored this phenotypic issue by measuring the expression of various genes involved in EMT by qRT-PCR in cells infected for 72 hours, 14 days, and 30 days. Specifically, we observed a significant decrease in E-cadherin expression starting from day 14 post-infection. Similarly, the expression of claudin-1, an epithelial marker, also decreased, though to a lesser extent, by day 30 post-infection (**Figure 3C)**. Conversely, two mesenchymal markers, fibronectin and TGFβ had an expression significantly increased at day 30 post-infection. (**Figure 3D)**. This led us to suggest that RPE cells persistently infected with ZIKV may have gradually lost their epithelial markers and begun an EMT.

## 4 Discussion

The possibility that certain pathophysiological processes following infection with pathogenic arboviruses may be due to long lasting and incomplete resolution of the initial infection is a growing documented hypothesis. Complete viral clearance failure is corroborated by the detection of viruses long after the symptomatic phase of infection. Several prospective studies conducted on cohorts of patients infected by ZIKV have revealed exceptionally long detection times for the virus in body fluids (up to 2 months in urine and nearly 6 months in semen), demonstrating the likely persistence of the virus in immunoprivileged niches and the potential for recurrence of ZIKV infection [33]. Viral persistence in the urogenital tract is considered to be responsible for the sexual transmission observed during recent Zika epidemics, leading the WHO to recommend the use of mechanical contraceptives during sexual intercourses, up to 3 months after known or presumptive infection. Prolonged viremia for around 20 days has also been observed in the case of dengue fever, with viral RNA detected in urine for up to 30 days from symptom onset [34]. The persistence of viruses in several organs long after the acute phase of the infection and blood virus clearance, is supported by the growing number of reported cases of donor-derived arboviral infections in organ transplant recipients [35]. These findings have led transplant units in endemic and epidemic areas to implement additional measures to detect the presence of various arboviruses in donors. This is the case for DENV, which is now screened in the donor’s urine and blood, prior to a kidney transplant [36]. In addition, the existence of viral reservoirs could explain late or long-term pathophysiological processes. CHIKV, an arbovirus member of the Alphavirus genus, causes chronic arthralgia that may be related to its persistence in the joints, which fuels a long-lasting and deleterious inflammatory environment [37]. Concerning ZIKV, regardless of the CZS observed at birth, there have been reports of neuropsychomotor developmental abnormalities occurring late in children who were born without symptoms but whose mothers were infected during pregnancy [38]. This suggests that ZIKV could persist for long periods in infected mothers, their fetuses, and newborns, with a possibility of delayed pathophysiological effects. Regarding ophthalmic manifestations associated with ZIKV infection, it should be noted that these are mainly reported subsequent to the acute phase. Several case reports document the first sign of uveitis or retinopathy 8 to 10 days after the onset of systemic symptoms, with detection of ZIKV by qPRT PCR in aqueous humor or conjunctival swabs [11,39,40]. These cases and studies in animal models suggest that tears and conjunctival fluid, as well as retinal cells, may constitute reservoirs for ZIKV. In addition, mouse models have allowed researchers to describe the eye damage induced by infection, revealing a disruption of the retinal layer that can persist for up to 60 days [14]. With our in vitro study on hTERT-RPE 1 cells, we provide additional data supporting the hypothesis of ZIKV persistence, suggesting that retinal pigment epithelial cells could be the primary infection point during acute retina infection, and the cell reservoir in the chronic forms of ophthalmic affections. We thus have shown that the infection is maintained for 30 days in RPE cells with regular production of infectious viral particles and infection rates oscillating between 10 and 25% during the timeline (**Figure 1 and 2**).

Upon infection, programmed cell death is one of the essential responses implemented by cells to eliminate an infectious agent. Its failure results in incomplete elimination of infected cells and the possibility of virus immune escape and persistence [41,42]. In the case of ZIKV, our team previously showed that in vitro infection of A549 cells elicited limited and delayed cytopathic effects. In addition, we demonstrated that ZIKV was able to control apoptosis in A549 cells by regulating Bcl2 family proteins and inhibiting CHOP-dependent UPR-induced death [43,44]. A review of the literature confirmed that the regulation of apoptosis was a key determinant of the pathological mechanisms, persistence capacity and transmission modes of ZIKV [45]. In our new study characterizing hTERT RPE cell infection, analysis of virus-induced cell death (**Figure 1**) showed similar results to those observed with the A549 cell line over a 3-day infection time course [46]. The rate of apoptotic cells then remains stable at around 15%-20% for cells infected for 6, 12 and 30 days (**Figure 2B**). Despite persistent basal infection, RPE cells cultured for 30 days appear to maintain a balance between cell death and cell growth that ensures renewal of the cell layer. It should be noted that the control-uninfected culture was also maintained for 30 days, with RPE entering quiescence, as described by Wang et al. [27]. To test the effect of this quiescent state on infection dynamics, we decided to proceed, after three weeks of culture, with trypsinization and reseeding of the chronically infected RPE cells. We observed that reactivation of the cell cycle led to a significant increase in the rates of infected cells and viral progeny that were measured ten days after passage and compared to non-passaged cells (**Figures 2C and 2D**). It appears that proliferating cells become more permissive. This phenomenon may result from shifts in cellular metabolism. It is now well established that the susceptibility of cells to viruses and the fate of viruses and infected cells are closely linked to the cell metabolic status. In a quiescent state, cells rely primarily on carbohydrate-derived substrates that enter the tricarboxylic acid (TCA) cycle, producing ATP via oxidative phosphorylation (OXPHOS) to maintain essential functions [47]. In contrast, proliferating cells favor aerobic glycolysis to support rapid biomass synthesis [48]. Cell passage initiates cell cycle re-entry and stimulates a metabolic shift toward glycolysis. This shift may create a favorable environment for viral replication, known to be highly biomass consuming, and account for the increased infection rate post-passage. Indeed, in our previous studies, we identified that OXPHOS promoted a more efficient antiviral response [49], and that a metabolic shift promoting mitochondrial oxidation of fatty acids limited ZIKV infection [50]. It should be noted that a growing body of literature supports that basal cellular metabolism (OXPHOS versus aerobic glycolysis), which is largely influenced by the growing or quiescent state, and its reorientation are important parameters for infection. They are thought to influence tropism, the efficiency of virus replication, and innate immune responses [51]. Moreover, it is also possible that renewal of the culture medium nullified the protection provided by soluble antiviral factors such as interferon, thus reigniting viral production. Previous studies have shown that ZIKV infection in RPE cells can elicit a strong antiviral response. Research conducted in primary RPE cells demonstrated that ZIKV promotes interferon signaling and upregulating a panel of antiviral molecules such as MX1, ISG15, OAS2, and CXCL10 [25]. Simonin et al. also investigated ZIKV infection in RPE derived from induced pluripotent stem cells (iPSCs). By 7 days post-infection, they observed antiviral and inflammatory responses in these RPE cells, mainly when the infection was carried out with African strains of ZIKV [26]. However, numerous studies report that the Asian strain of ZIKV, responsible for the last outbreaks, has the ability to limit type I IFN responses, promoting a rather low-noise infection (e.g. in human neural stem cells) that could allow the virus to persist in the host and cause long-term defects [52–54]. We need to confirm in our model whether the regulation of the innate immune response contributes to the inability of the RPE to clear the virus. We hypothesize that a combination involving apoptotic restriction, cell cycle stage, and modulation of the innate immune response would likely facilitate ZIKV persistence in RPE cells.

Finally, the most striking observation of our study was that prolonged infection of hTERT RPE-1 cells with ZIKV led to a morphological transition together with a loss of epithelial characteristics (**Figure3**). When we noticed that the cells took on an elongated, spindle-shaped appearance typical of fibroblasts, we hypothesized that an epithelial-mesenchymal transition (EMT) was occurring. Under our conditions, only two mesenchymal markers were found to be elevated 30 days post-infection (fibronectin and TGF-β). However, since the 14th d.p.i., we have observed a downward regulation of epithelial markers such as claudin-1 and E-cadherin. This suggests that the cells lost some of their properties related to the expression of genes encoding proteins required in epithelial cell-cell junctions. Claudins are tight junction components, essential for the cohesion of the RPE monolayer and its impermeable properties [55]. A loss of integrity in tight junctions results in many eye diseases. Knock-out mice for the gene encoding claudin-1 present an hyperpigmentation of the retinal pigment epithelium, and lipid and drusen-like deposits similar to the clinical signs observed in humans with age-related macular degeneration [56]. Cadherins are a family of membrane proteins involved in calcium-dependent intercellular adhesion and E-cadherin plays a crucial role in the formation of adherens junctions between cells in the blood-retinal barrier. Mutations in genes encoding cadherins have been identified as causes of hereditary retinal degeneration [57]. Knockdown of E-cadherin was also shown to affect not only adherens junctions but also tight junctions, compromising epithelial function [58]. The decrease in the expression of these genes in our chronically infected RPE cell cultures is therefore a strong indicator of a probable loss of epithelial properties. Additional experiments evaluating the integrity of the RPE cell layer, such as measurements of trans-epithelial electrical resistance or measurements of dextran permeability in transwell cultures, are needed. It can already be assumed that a dysfunction of the blood-retinal barrier could be a consequence of this loss of epithelial markers associated with the locally persistence of ZIKV. An interesting aspect of our observations is that all the RPE cells, not just the 10% infected cells, are affected by the morphological change. This change could therefore be driven by a soluble factor. It is known that EMT is mediated by TGF-β signaling. In the case of proliferative vitreoretinopathy, an EMT of retinal pigment epithelium cells triggers the pathophysiology. During the process RPE cells differentiated into myofibroblasts in the presence of TGF-β, and acquired migration, proliferation, and contraction capabilities that resulted in fibrotic complications leading to blindness [59]. In our study, we found that cells infected with ZIKV for 30 days displayed a nearly fourfold increase in the expression of genes encoding fibronectin and TGF-β. We would, of course, need to quantify the TGF-β secreted in the media to verify its role in the phenotypical change of RPE and the gain of fibrotic characteristics.

In conclusion, disruption of the BRB epithelial phenotype could be a step of the viral infection time course, thus providing an easy access to an immuno-privileged viral niche, from which ZIKV could chronically persist. This pattern of phenotypic change in the RPE, with loss of epithelial properties and gain of fibrotic mesenchymal features is observed in several eye diseases. It would account for a pathological process induced by ZIKV persistence and explain the late ophthalmic complications observed in cases of retinopathy and visual impairments following infection. Taken together, our results shed a new light on a previously uncharacterized infectious process, and position RPE cells as interesting targets in the treatment of persistent arboviral infections.

## Author Contributions

Conceptualization, D.E.S., A.M., G.L., W.V., and P.K-T.; Methodology; resources; writing—original draft preparation, D.E.S., A.M., W.V., and P.K-T.; Writing—review and editing, P.K.-T.; Visualization, P.K.-T. and W.V.; Supervision, P.K.-T.; All authors have read and agreed to this version of the manuscript.

